# Patient-specific logic models of signaling pathways from screenings on cancer biopsies to prioritize personalized combination therapies

**DOI:** 10.1101/422998

**Authors:** Federica Eduati, Patricia Jaaks, Christoph A. Merten, Mathew J. Garnett, Julio Saez- Rodriguez

**Affiliations:** European Molecular Biology Laboratory (EMBL), Genome Biology Unit, Meyerhofstrasse 1, Heidelberg, Germany; European Molecular Biology Laboratory, European Bioinformatics Institute (EMBL-EBI), Wellcome Trust Genome Campus, Hinxton, UK; Joint Research Centre for Computational Biomedicine (JRC-COMBINE), RWTH Aachen University, Faculty of Medicine, Aachen, Germany; Dept. Biomedical Engineering, Eindhoven University of Technology, Eindhoven, The Netherlands; Wellcome Trust Sanger Institute, Wellcome Trust Genome Campus, Hinxton, Cambridgeshire, United Kingdom; Institute for Computational Biomedicine, Heidelberg University, Faculty of Medicine, BIOQUANT-Center, 69120 Heidelberg, Germany.

## Abstract

Mechanistic modeling of signaling pathways mediating patient-specific response to therapy can help to unveil resistance mechanisms and improve therapeutic strategies. Yet, creating such models for patients, in particular for solid malignancies, is challenging. A major hurdle to build these models is the limited material available, that precludes the generation of large-scale perturbation data. Here, we present an approach that couples *ex vivo* high-throughput screenings of cancer biopsies using microfluidics with logic-based modeling to generate patient-specific dynamic models of extrinsic and intrinsic apoptosis signaling pathways. We used the resulting models to investigate heterogeneity in pancreatic cancer patients, showing dissimilarities especially in the PI3K-Akt pathway. Variation in model parameters reflected well the different tumor stages. Finally, we used our dynamic models to efficaciously predict new personalized combinatorial treatments. Our results suggest our combination of microfluidic experiments and mathematical model can be a novel tool toward cancer precision medicine.

## Introduction

Charting the dynamic wiring of signaling networks is of paramount importance to understand how cells respond to their environment. Identifying the differences in this wiring between normal and cancerous cells can shed light on the pathophysiology of tumors and pave the way for novel therapies (Saez-Rodriguez *et al*, 2015; Werner *et al*, 2014; Zañudo *et al*, 2018). A powerful tool to gain insight into these processes is to monitor the response of cells to multiple perturbations. When combined with mathematical modeling, such data can be used to determine cell type-specific wiring phenomena, predict efficacy of drug treatments, and understand resistance mechanisms (Eduati *et al*, 2017; Hill *et al*, 2017; Saez-Rodriguez *et al*, 2011; Merkle *et al*, 2016; Klinger *et al*, 2013).

Application of this strategy has been limited so far to *in vitro* contexts, as the experimental technologies to generate perturbation data require large amounts of material, which are unavailable from most primary tissues such as solid tumors. Recently developed organoid technologies allow to generate large amounts of material *ex vivo*, enabling such screens in principle. However, they would be associated with large costs and, while recapitulating some of the features of the tumor physiology, the cells unavoidably diverge from the primary tumor as they are grown *ex vivo (Letai, 2017)*. We have recently developed a novel strategy based on microfluidics that enables testing apoptosis induction upon a good number of conditions (56 with the current settings, with at least 20 replicates each) starting from as little as one million viable cells. Cells are encapsulated in 0.5 μl plugs together with an apoptosis assay and single or combined drugs. Using valves to control individual fluid inlets allows the automatic generation of plugs with different composition. These Plug-Based Screenings (PBS) are suitable to collect such drug response datasets even with the very limited number of cells available from tumor resection biopsies (Eduati *et al*, 2018).

In this study, we set out to construct cell type-and patient-specific models from the drug response data obtained using the PBS technology (overview of the pipeline in **Figure 1A**). We took advantage of our tool CellNOptR (Terfve *et al*, 2012), to train a general network of the underlying intrinsic and extrinsic apoptosis pathways from data obtained for two cell lines and biopsies from four pancreatic tumor patients at different stages. We found the models to be a useful tool to understand specific pathway deregulations and to predict new patient-specific therapies.

**Figure 1.**
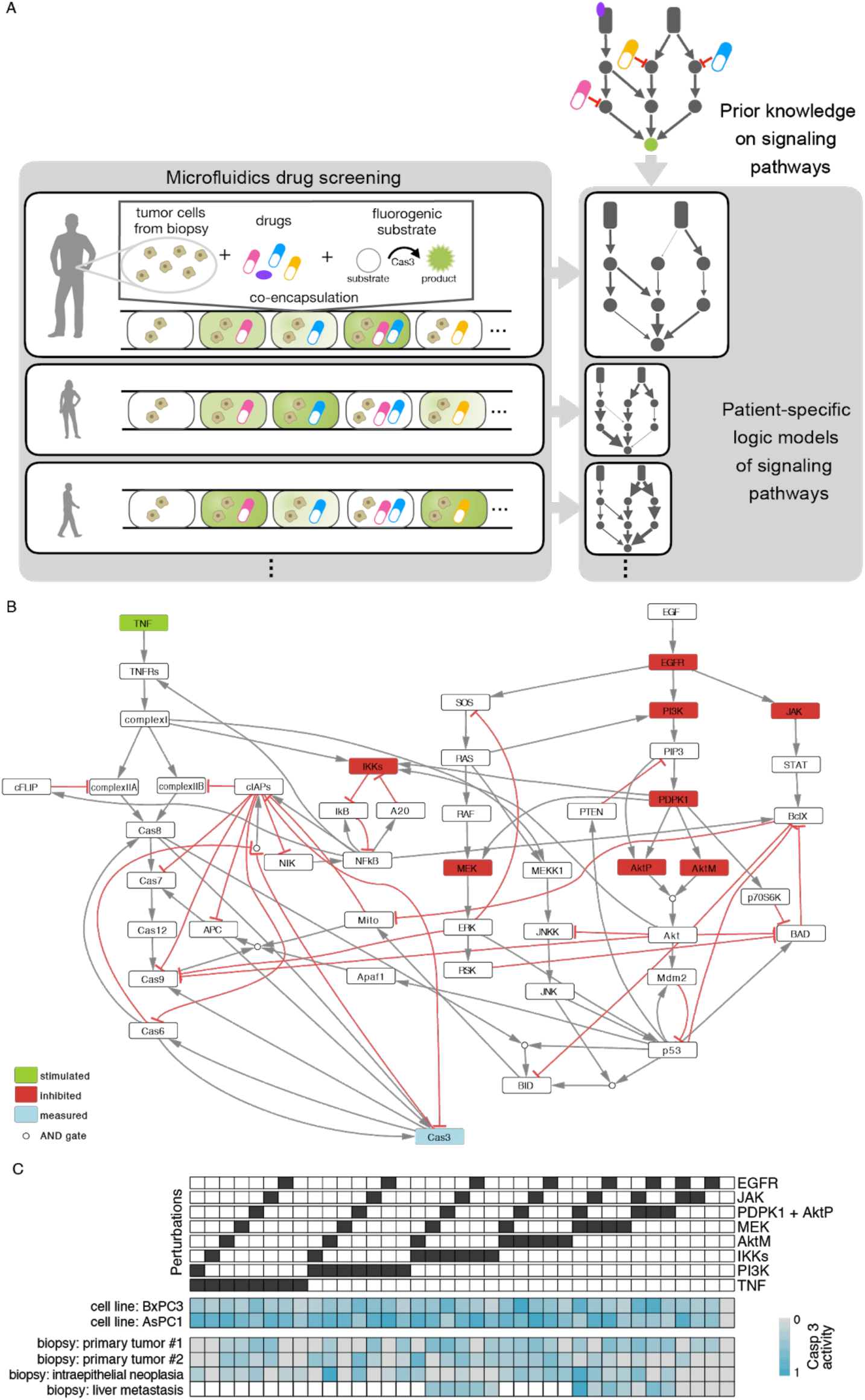
Apoptosis pathways and experimental data. A. Overview of the pipeline: Patient-specific mathematical models are built for each patient from the combinatorial microfluidics plug based screening (PBS) data measured on live cells from cancer patient biopsies. B. Logic model of intrinsic and extrinsic apoptosis pathways regulating Cas3 (our readout, blue node), including all nodes which are perturbed (stimulated in green, inhibited in red) in the experiments. C. Experimental data consisting of 37 different experimental conditions (columns) for 6 samples (rows; 2 cell lines and 4 biopsies).

## Results

### Data and modeling of apoptosis pathways

Experimental data were generated using a novel plug-based microfluidics screening (PBS) platform, which allowed performing combinatorial drug screening of biopsies from human tumors (Eduati *et al*, 2018) (see **Methods**). Data represents caspase-3 (Cas3 in **Figure 1B**, marked in blue) activation after perturbation with 10 different compounds including seven kinase inhibitors (targeting IKKs, MEK, JAK, PI3K, EGFR, AKT, PDPK1 -inhibited nodes are depicted in red in **Figure 1B**), one cytokine (TNF, stimulated node, in green) and two chemotherapeutic drugs (Gemcitabine and Oxaliplatin). All 10 compounds were tested alone and in all 45 possible pairwise combinations (**Figure 1C**) on two pancreatic cancer cell lines (AsCP1 and BxPC3) and biopsies from four patients with pancreatic tumors at different stages (one intraepithelial neoplasia, two primary tumors and one liver metastasis).

To investigate the signaling mechanisms behind the differential drug responses of our cell lines and patients, we derived a general logic model of apoptosis pathways involved in the regulation of Cas3 (our measurement), which is considered as effector node and indicator of apoptosis. Models were then trained using the patient-specific experimental data to obtain personalized models. The general model (**Figure 1B**) was built integrating information derived from literature and from public repositories (details in **Methods** section). The model describes both intrinsic (mediated by the mitochondria, named Mito in the model) and extrinsic (mediated by Tumor Necrosis Factor Receptors TNFRs) apoptotic signals, including nodes encoding for both anti-and pro-apoptotic effects. We incorporated in the model all nodes perturbed by specific compounds in our screening such as targeted drugs (kinase specific inhibitors) and the cytokine TNF. The effect of chemotherapeutic DNA damaging drugs was not included in the model since they inhibit DNA replication rather than acting directly on specific signaling nodes. Since our screening included two AKT inhibitors (i.e. MK-2206 and PHT-427) with different mechanisms of action (allosteric and PH domain inhibitors, respectively), they were modeled as acting on two different nodes (AktM and AktP respectively), both needed for the activation of AKT.

The logic model includes AND gates (dots in **Figure 1B**) when all upstream regulators are needed to activate a node, while cases with multiple independent regulators are considered as OR gates. The logic model is interpreted using the logic based ordinary differential equation formalism (logic ODEs) (Wittmann *et al*, 2009) as implemented in CellNOptR (Terfve *et al*, 2012). This formalism allows to maintain the simple causal structure of logic models, while considering also the dynamic nature of the interactions and the continuous scale for the activation of the nodes, by using ODEs. As previously described (Eduati *et al*, 2017), we consider one parameter for each edge *j → i* in the network, which characterize the strength of the regulation of species *i* dependent on species *j* and one parameter for each node *i*, which represents the responsiveness of the node (see **Methods**).

### Calibration of the apoptosis model for cell lines

The parameters of the generic model were fitted separately to the data of each cell line, resulting in specific models tailored to the experimental data for each cell line (more details in **Methods** section). Parameter fitting was repeated 10 times and performances were assessed using different metrics to compare model simulations with the experimental data, showing good and quite robust performances (average metrics for AsPC1 and BxPC3 respectively: Pearson correlation 0.72, 0.74; mean squared error 0.03, 0.02; coefficient of determination 0.5, 0.5; **Appendix Figure S1A)**. Model simulations for the best specific models were compared with the corresponding measured experimental data, showing a very good agreement (Pearson correlation equal to 0.89 and 0.83 for AsPC1 and BxPC3, respectively; **Appendix Figure S1B-C**).

The calibrated models for these two cell lines were then used to uncover potential differentially regulated mechanisms which are behind the different drug responses of the cell lines. Due to the limited number of data and the complex nature of the signaling pathways involved in activation of apoptosis, not all model parameters can be estimated with the same confidence. In order to estimate the variability of the optimised parameter values, we derived a bootstrapped distribution for each parameter for each cell line, by repeating the optimisation 500 times while randomly resampling the data with replacement. These distributions were then used to compare the two cell lines using statistical tests to highlight significant differences (Wilcoxon sum rank test, adjusted p-value < 0.01, effect size > 0.2, see **Methods**) as represented in **Figure 2A**. The comparison revealed some regulatory mechanisms which are upregulated in either AsPC1 or BxPC3. Additionally, the dynamic of Cas3 activation appears to be faster in AsPC1 (node border in green). Main differences involve the PI3K-Akt pathway. The main linear pathway is more active in BxPC3, whereby the negative feedback loop, from p53 to PIP3 mediated by PTEN, is stronger in AsPC1. These differences in the model parameters cause changes in the dynamic behaviour of the system (**Figure 2B**) and are behind the differential activation of Cas3 in response to drugs.

**Figure 2.**
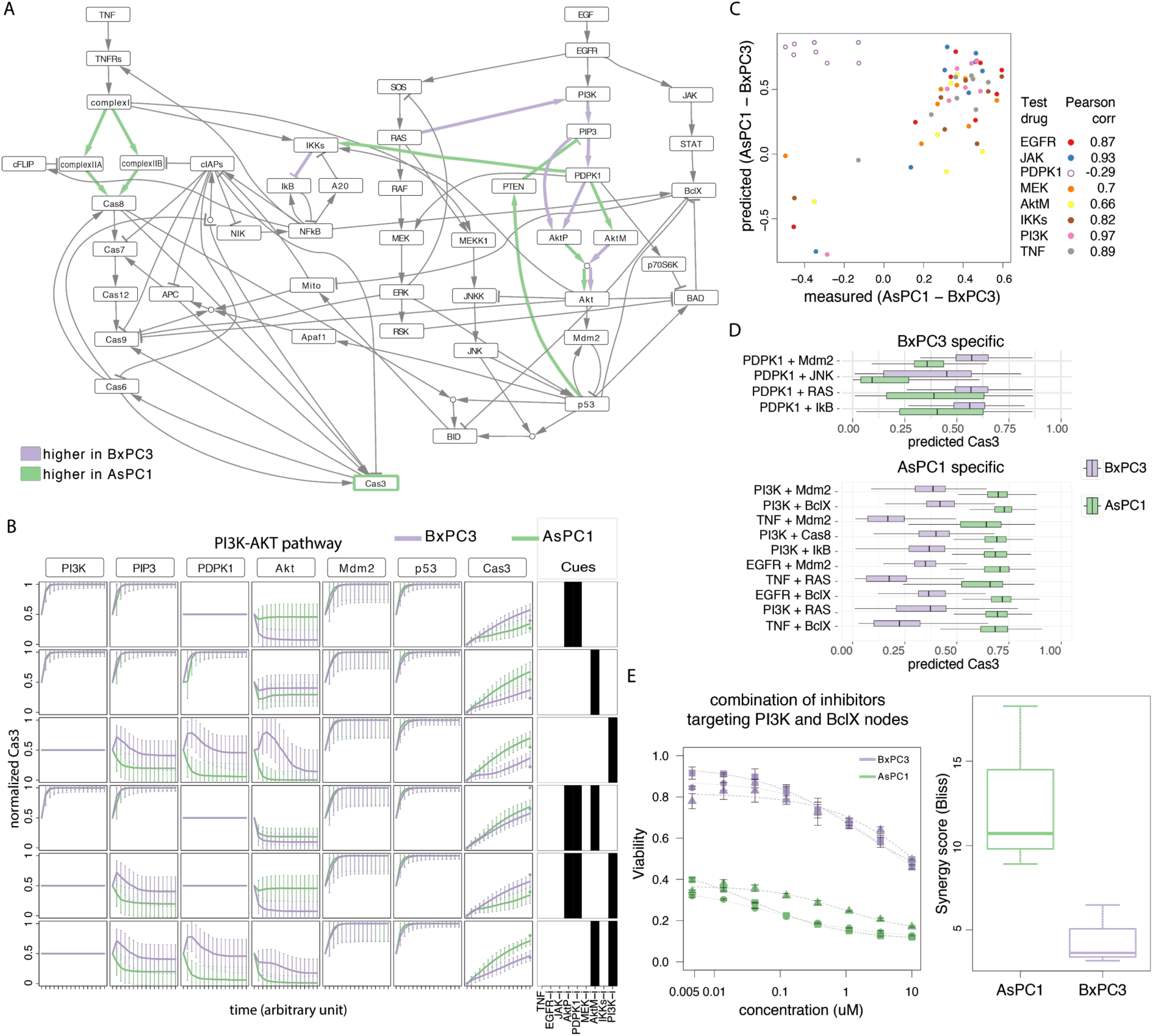
Models, predictions and validation for AsPC1 and BxPC3 cell lines. A. Differentially regulated mechanisms in AsPC1 or BxPC3, highlighted in green or purple depending on whether the corresponding estimated parameters are higher in AsPC1 or BxPC3, respectively. B. Time course simulation of PI3K-AKT pathway and related inhibitors in AsPC1 and BxPC3. Lines represent median values and error bars represent standard deviation from the bootstrapped simulations. C. Assessment of model predictions by cross-validation, removing for each repetition all experiments involving one of the 8 drugs (the corresponding drug targets are reported in the legend) from the training set and using it as test set. Bad predictions (Pearson Correlation < 0.6 -which corresponds to removal of drug targeting PDPK1) are marked with empty dots. D. New drug combinations predicted to be highly specific for each cell line (only top 10). E. Experimental validation of the combination of PI3K (Taselisib) and BclX (Navitoclax) inhibitors predicted to be specific for AsPC1. Data shown are for three biological replicates, with 3 technical replicates each, for Navitoclax at 2.5 uM and different concentrations of Taselisib (8 points 1:3 dilution series), complete data are shown in Appendix. Boxplot (right panel) shows the corresponding synergy scores (Bliss model).

We then investigated if these differences in dynamic behaviour could be derived from their genetic makeup (Garcia-Alonso *et al*, 2018). From the proteins in our model, only KRAS is functionally mutated in AsPC1 and TP53 in BxPC3. Furthermore, no known direct regulators of the nodes in our model - from those in Omnipath (Türei *et al*, 2016), a compendium of pathway resources - was mutated. Interestingly, KRAS and TP53 are indeed involved in the pathways that we found to be differentially activated between the two cell lines, but on their own they cannot explain the differences in pathway structure in terms of strength of regulations. Therefore, information on mutations alone would not be sufficient to describe the dynamics of the pathways that mediate apoptosis upon drug treatment. The same holds when looking at basal transcriptomics (Iorio *et al*, 2016) (**Appendix Figure S2**), supporting the observation that static data are not sufficient to investigate the dynamics of a complex system.

### Model predictions and validation

We decided to test the predictive power of our cell line-specific mathematical models in two ways: 1) using cross-validation on the existing dataset, and 2) predicting the effect of new drug combinations that can be experimentally tested.

First, optimisation was repeated randomly selecting eight conditions as validation set and using the remaining 36 for training (bootstrapping 100 times). This procedure was repeated for 20 randomised test sets and results show a good correlation between the predictions and the measurements in the validation set (average Pearson Corr = 0.7). Even when the cross-validation was repeated removing all the experiments involving a specific drug each time (instead of random ones, **Figure 2C**), predictions are still very good (Pearson Corr range 0.66-0.97) for all drugs except PHT-427 (Akt and PDPK1 inhibitor, Pearson Corr = -0.29). This implies that experiments with PHT-427 are essential to define the models.

Mathematical models were then used to simulate the effect of different drug combinations acting on the pathways that were not previously tested using PBS. For each cell line we simulated the effect of 186 new perturbations (12 single drugs and 162 drug combinations), by inhibiting the corresponding node in the model. Varying confidence of model parameter estimation from the available data is expected to affect the ability to predict certain conditions. By using the family of models optimised using bootstrap, we obtained a distribution of the predicted activation of Cas3 in response to the different simulated conditions, therefore retaining information on the confidence we have for each prediction. In particular we focused on the predictions which were significantly different between the two cell lines (Wilcoxon sum rank test, adjusted p-value <0.01, effect size > 0.3, see **Methods**). To select drug perturbations highly specific for each cell line, we considered only those with median predicted value for Cas3 > 0.45 and ranked them based on the effect size. This results in 4 conditions specific for BxPC3 and 49 for AsPC1 (**Figure 2D**, **Appendix Figure S3**).

We then tested experimentally one of the top combinatorial therapies predicted to be highly specific for AsPC1 based on our mathematical models, consisting in the combination of a PI3K inhibitor (Taselisib) with a drug targeting the BclX node (Navitoclax, a Bcl-2/Bcl-xL/Bcl-W antagonist). Agents targeting the PI3K pathway in combination with a Bcl-2 family inhibitor have been previously suggested to be relevant in the context of pancreatic cancer (Tan *et al*, 2013). Hence, being able to predict the efficacy of this combination for specific patients (or cell lines in this case) would be highly desirable. Our validation experiments proved that the combination of Taselisib and Navitoclax is more efficacious and synergistic (based on Bliss independence model) in AsPC1 than in BxPC3 cells (**Figure 2E, Appendix Figure S4**), confirming our model-based predictions.

### Personalized apoptosis models for patients’ tumors

The same fitting pipeline previously described for the cell lines was applied to the data from the four pancreatic tumor biopsies (intraepithelial neoplasia, two primary tumors and one liver metastasis) to obtain personalized models. Patient-specific parameter distributions were used to investigate patient heterogeneity at the level of mechanisms involved in apoptosis signaling pathways. Results are summarized in **Figure 3A**, showing the 16 (out of 93) parameters which are different in at least one patient (Kruskal-Wallis rank sum test, adjusted p-value < 0.01, effect size > 0.2, see **Methods**). For these parameters, we also performed post-hoc pairwise statistical tests to directly compare all patients (Wilcoxon sum rank test, adjusted p-value < 0.01, effect size > 0.2, see **Methods**). For instance, for the parameter representing the EGFR → JAK regulation the null hypothesis of equal distribution is not rejected when comparing the two primary tumors (lower two boxes in gold) between themself and with respect to intraepithelial neoplasia (top-left box, half gold next to the corresponding interaction in **Figure 3A**), however it is rejected when comparing each of them with liver metastasis (top-right box, in cyan). Also the comparison of intraepithelial neoplasia and liver metastasis suggests that the two samples do not come from different distributions (top boxes, both cyan).

**Figure 3.**
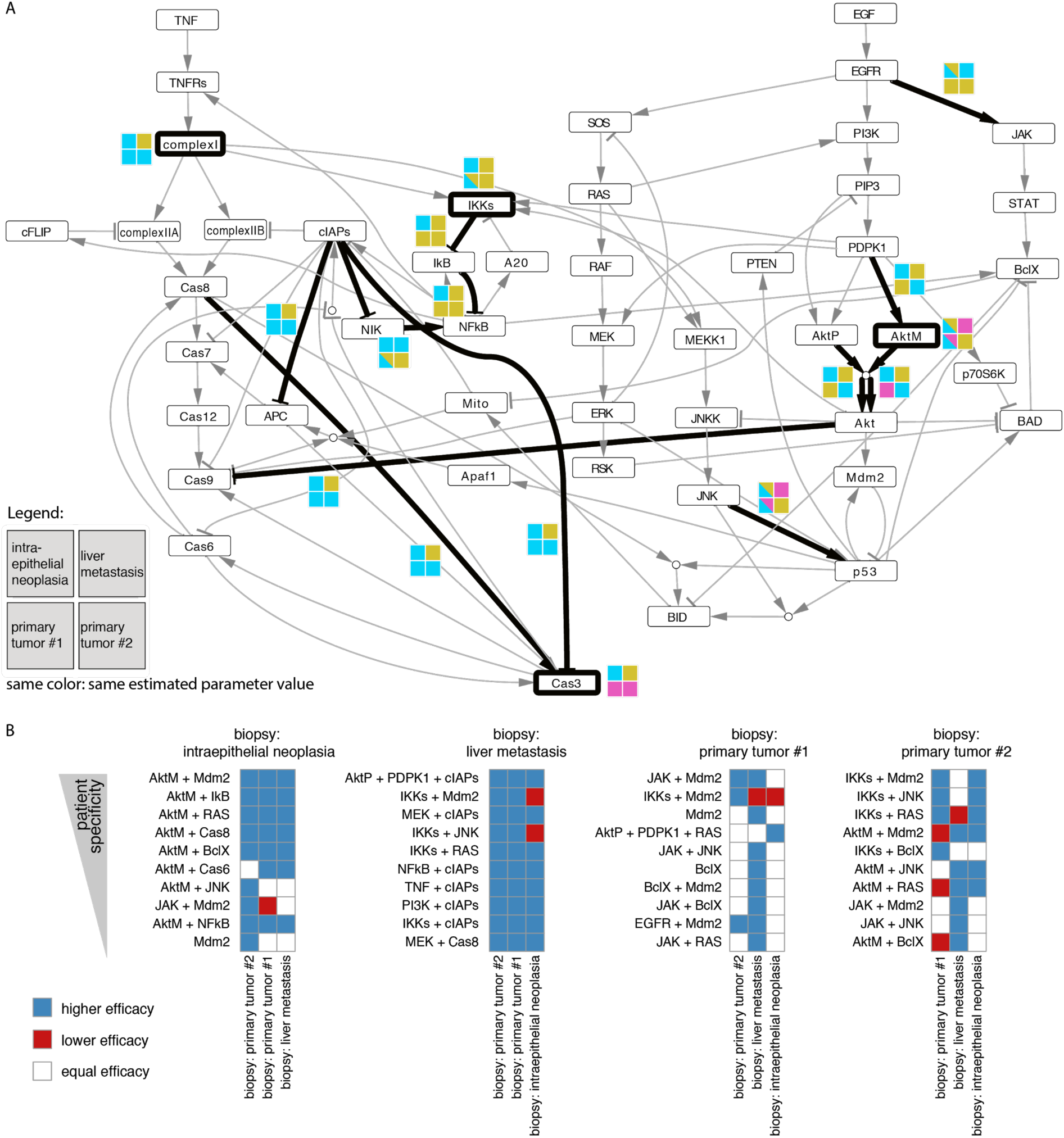
Patient-specific models of signaling pathway. A. Mechanisms which are differentially regulated among the patients are highlighted with thick black lines. Colored squares represent the same distribution (same color) or differential distribution (different colors) across the four patient samples. B. Patient-specific predictions of new drug combinations. Predictions are ranked for each patient and color coded to compare the efficacy with the other patients.

Overall, the most different sample is the liver metastasis (different from all the others in 7 of the 16 heterogeneous parameters (44%)), especially in the extrinsic apoptosis pathway mediated by complex I, cIAPs and Cas8. This larger dissimilarity could be justified by the difference both in stage and in tissue, since all other samples were resected from the pancreas. Also, the intraepithelial neoplasia shows a quite high level of dissimilarity (37.5%), localized in particular in the IKKs-NFkB pathway, which could reflect the more advanced stage of the disease. Interestingly, the two primary tumors are the most similar with each other (similar in 11 out of the 16 parameters, corresponding to 69%). However, significant differences were found especially in the PI3K-AKT pathway, similarly to what we observed for the two pancreatic cancer cell lines. Importantly, these similarities, which reflect the different tumor stages, were not evident directly from the data, where primary tumor #1 clusters closer to the liver metastasis and primary tumor #2 closer to the intraepithelial neoplasia (**Appendix Figure S5**).

### Model-based prioritisation of new personalized treatments

The personalized models can be used not only to investigate the patient-specific deregulated mechanisms, but also to predict novel experimental conditions as previously shown for the cell lines. For example, we can simulate the effect of a new kinase inhibitor by inhibiting the corresponding node in the model and predict its effect on Cas3 and thus on apoptosis. By implementing *in silico* testing, we can increase the throughput of our screening method for each patient, allowing to predict the effect of new potential therapies which cannot be experimentally tested due to limited biopsy material.

For each patient, we simulated the effect of 12 new single and 162 combinatorial therapies (186 *in silico* perturbations in total) targeting nodes in our model. Having applied bootstrap when deriving our personalized optimized models, as previously described, we obtained a distribution of the predictions for each simulated perturbation. This allowed us to perform statistical tests to compare the effect of the same treatment across patients and focus on patient-specific effects, removing 129 out of the 186 treatments that were not statistically different between patients (Kruskal-Wallis rank sum test, adjusted p-value < 0.01, effect size > 0.3, see **Methods**). With this step, we removed treatments that could not be predicted with sufficient confidence for any patient (broad distributions), and the treatments showing high predicted efficacy for all patients, that could be likely due to general toxicity. For the remaining 57 conditions, we performed also post-hoc comparison between all patient pairs (Wilcoxon sum rank test, adjusted p-value < 0.01, effect size > 0.3, see **Methods**). Similar to what we noticed comparing model parameters, using these predictions we observed that primary tumors behave most similar as they show no statistical difference in 79% of the cases. This is much higher than the similarity observed in comparison with intraepithelial neoplasia (60% and 43% for primary tumor #1 and #2, respectively), or the liver metastasis (12% and 11% respectively for primary tumor #1 and #2).

Finally, in order to prioritize new patient-specific promising treatments, we ranked the simulated perturbations for each patient by patient specificity (i.e. higher effect size in the pairwise comparison); **Figure 3B** shows the top ten for each patient. Interestingly, there are three treatments showing strong potential (among the top ten) for both primary tumors. Two of these consist in targeting Mdm2 in combination with JAK and IKKs, respectively. Mdm2-p53 binding is known to be an important target in pancreatic cancer, where TP53 mutations occur in 50-70% of the patients (Morton *et al*, 2010). Finding treatments to combine with drugs disrupting this binding, like Nutlin-3 (Khoo *et al*, 2014), is currently of great interest (Bykov *et al*, 2018) especially in pancreatic cancer (Izetti *et al*, 2014). In particular, activation of the JAK-STAT pathway has been shown to be common in pancreatic cancer (Matsuoka & Yashiro, 2016) and is often associated with TP53 mutation (Wörmann *et al*, 2016), suggesting that targeting Mdm2-p53 and JAK could be indeed a promising combination therapy for some patients. Based on our predictions, combinatorial targeting of Mdm2 and JAK is also efficacious in the intraepithelial neoplasia, while targeting Mdm2 in combination with IKKs is more efficacious in the liver metastasis.

## Discussion

Given the intrinsic complexity of cancer, experimental data obtained recording the cellular response to perturbations are essential to study cancer cells as a dynamic system. This functional data provides complementary information to that obtained by genomic profiling in steady state, and is particularly relevant for investigating therapeutic efficacy of anticancer drugs (Letai, 2017; Yaffe, 2013). Further knowledge can be extracted by analysis of this experimental perturbation data via mathematical modeling, providing a rationale for mechanism-based interpretation of drug response.

We here present an approach to effectively build mechanistic models from integration of large-scale perturbation datasets and prior knowledge of the underlying pathways. Because of the low material needed by our recently developed PBS platform (Eduati *et al*, 2018), our approach can be applied not only to *in vitro* but also to *ex vivo* settings, as demonstrated in this study. Our tool allows us to dissect functional differences in the signaling pathways by comparing the model parameters. These parameters recapitulate similarity between different tumor stages better than the drug screening data, suggesting that they can shed light on the molecular basis of tumors at the individual patient level. In addition, the tool can be used to rationally select efficacious combination therapies, as illustrated on the study in cell lines. We chose a simple logic formalism so that we could efficiently model large networks despite measuring a single readout. *In silico* and *in vitro* analyses demonstrate that our approach is robust. In summary, our combination of mathematical modeling and *ex vivo* perturbation data helps to investigate possibly deregulated mechanisms (or pathways) and to explain specific responses to drugs directly on patients’ biopsies.

The PBS platform can be extended in the future to provide richer data, and thereby improve the mathematical models. We have so far generated PBS data with a large number of perturbations but a single readout (apoptotic marker). While informative, in particular to study the effect of anti-cancer drugs, it is limited to capturing a subset of signaling networks. PBS can in principle be applied to other markers, as well as connected to richer output technologies, such as high-content imaging, or single-cell RNA sequencing. We expect that the breadth and depth of our models will increase when expanding readouts. In addition, our modeling approach is not limited to PBS, but can be used with different types of data describing molecular or phenotypic changes upon perturbation. Besides the readouts, extension to multi-time-point measurements will provide additional insight into the dynamics and feedback regulation of the system. Finally, a higher granularity in drug concentrations tested will provide information on intermediate effects. We expect that further developments of technologies for functional screening of cancer patient biopsies will follow in the near feature (Letai, 2017), and this will reflect in further improvement of patient-specific mathematical models that can be obtained using our pipeline.

Considering a family of models allows us to account for cell signaling heterogeneity for both estimated parameters values and model predictions (Kim *et al*, 2018). This is currently taken into account when comparing models and when making predictions of efficacious therapy. By using statistical tests, we consider as promising only combinations that are robustly more efficacious for each individual patient/cell line. Having few cells per plug (~100) and many replicates (at least 20 per condition), we have collected information on the heterogeneity of cellular response to drugs within a patient sample, which could be taken into account when building the model. Statistical models could be used in the future also to distinguish between the variability due to technical noise (same for all plugs) and the variability due to heterogeneity of cellular response to drugs (specific for each condition). Alternatively, different cell types can be sorted out prior to the PBS experiments, to obtain cell type-specific information.

Generation of perturbation data followed by mathematical modeling has proven to be a powerful tool to study cancer biology and therapies *in vitro*. The insights from *in vitro* models can be extrapolated to patient data using a patient’s static profiling, such as gene expression (Fey *et al*, 2015). When no other data is available, this is certainly a very valid strategy to generate personalized models. Our work shows however, that this basal information cannot recapitulate the insights obtained by data upon perturbation. If one can generate such data directly from patient samples, we should be able to generate more precise models that provide more accurate insights and predictions. We believe the strategy presented in this work can contribute to the development of functional precision cancer medicine.

## Methods

### Microfluidics setup, screened compounds and samples

Data were generated using our plug-based screening (PBS) platform as presented in (Eduati *et al*, 2018). A cell suspension is generated from cell lines in culture or from patient biopsies (**Figure 1A**). A microfluidics chip is then used to automatically generale plugs with different chemical composition, using valves that can be opened and closed using a Braille display. In each plug, cells (about 100) are encapsulated together with one or two compounds and a rhodamine 110 (green-fluorescent dye) based substrate of caspase-3 ((Z-DEVD)2-R110), which is a marker of apoptosis. The activation of caspase-3 causes the cleavage of the substrate and the subsequent release of the green fluorescence in the plug. Alexa fluor 594 (orange-fluorescent dye) is added to the cell suspension to verify the proper mixing of the different components in each plug.

Samples are produced in a sequential way in multiple replicates (12 for perturbations, 20 for untreated control), and each sample is followed by a corresponding barcoding sequence produced using two different concentrations of Cascade Blue dye (blue-fluorescent dye) to encode the sample number in binary digits. The full sequence of conditions is repeated at least twice (resulting in a total of at least 24 replicates per perturbation). Aqueous plugs are separated by mineral oil plugs to avoid cross-contamination. All plugs are collected in a tube and incubated overnight for 16 hours at 37°C and 5% carbon dioxide. Fluorescence in three channels (green, orange, blue) is measured for each plug by exciting it with lasers (375, 488 and 561 nm) and detecting the emissions with corresponding three photomultiplier tube (PMT) detectors (450, 521 and >580 nm).

The 10 screened compounds (alone and in all pairwise combinations) include two cytotoxic drugs (Gemcitabine, Oxaliplatin), standard-of-care for pancreatic cancer, 7 kinase inhibitors (ACHP: IKKi, AZD6244: MEKi, Cyt387: JAKi, GDC0941: PI3KI, Gefitinib: EGFRi, MK-2206: AKTi. PHT-427: AKTi & PDPK1i) and one cytokine (TNF).

In accordance with the Declaration of Helsinki of 1975, human pancreas biopsies (primary tissue samples) were obtained during routine clinical practice at University Hospital Aachen, Aachen, Germany, and were provided by the RWTH Aachen University Centralized Biomaterial Bank (cBMB) according to its regulations, following RWTH Aachen University, Medical Faculty Ethics Committee approval (decision EK 206/09). Sample processing at the EMBL in Heidelberg, Germany, was approved by the EMBL Bioethics Internal Advisory Committee.

### Building the apoptosis pathway model

The logic model shown in **Figure 1B** was derived by manual literature curation starting from the model described by Mai and Liu (Mai & Liu, 2009) and integrating additional information in order to include all nodes perturbed in our experiments and to well describe pathway cross-talks. Logic rules were also adopted from the Boolean model of apoptosis by (Schlatter *et al*, 2009). We modeled both the intrinsic (mediated by the mitochondria) and the extrinsic (mediated by death receptors, TNFRs) apoptosis signal including nodes encoding both anti-and pro-apoptotic effects. Binding of TNF to TNFRs activates the extrinsic pathway mediated by Caspase-8 (Cas8 in **Figure 1B**) activation of Caspase-3 (Cas3). The two distinct Caspase-8 activation pathways (Wang *et al*, 2008) are represented by the cascade involving complex I (composed of RIPK1, TRADD, TRAF2), which induces the formation of two different Caspase-8 activation complexes: complex IIA (TRADD, RIPK1, FADD, Pro-caspase 8) and complex IIB (RIPK1, TRADD, FADD, Pro-Caspase 8, cFLIP) that can be inhibited by cFLIP and cIAPs respectively. For simplicity, Caspase-8 is modeled as a separate node (Cas8) regulated by the two complexes. TNF can also regulate the intrinsic pathway through the activation of NFkB (anti-apoptotic node) by removal of its inhibitor IkB. The activation of the intrinsic pathway is executed by the mitochondria through the release of SMACs (second mitochondria-derived activator of caspases) and Cytochrome c. The former deactivates IAPs, which are anti-apoptotic proteins, the latter binds to Apaf1 (Apoptotic protease activating factor-1) and pro-caspase9 which is converted to its active form of Caspase-9 (Cas9) and in turn activates Caspase-3 (Cas3). Both Akt and ERK have an anti-apoptotic effect by phosphorylating BAD (Balmanno & Cook, 2009) and thus unbinding it from BclX and this can be modelled as an OR gate (She *et al*, 2005). We also included the pro-apoptotic effect of ERK as regulator of p53 (Cagnol & Chambard, 2010). Additional cross-talks from RAS to MEKK1 and PI3K pathways were included as described by Grieco and colleagues (Grieco *et al*, 2013). Additional interactions between nodes in the network were found using Omnipath (Türei *et al*, 2016) and through manually curating the literature supporting the interactions in the databases. For example, in this way we found support for potential context dependent cross-talks from PDPK1 to MEK (Aksamitiene *et al*, 2012; King *et al*, 2000; Borisov *et al*, 2009) and to IKK/NFkB signaling (Tanaka *et al*, 2005) which were therefore added to our prior knowledge network.

### Data normalisation and formal definition of logic ODEs

Data from the PBS screening were pre-processed using the pipeline for data analysis and quality assessment described in (Eduati *et al*, 2018) and implemented in R (https://github.com/saezlab/BraDiPluS). In short, we used the signal in the orange channel (see description of the Microfluidics setup) to discard corrupted data corresponding to improperly formed plugs. For each screened condition in each run (i.e. full sequence of all tested conditions - see description of the microfluidics setup), we computed the median across replicates (12 plugs produced per condition) and the corresponding z-score. Additionally we compute the FDR-corrected p-value with respect to the untreated control (one-sided Wilcoxon rank-sum test). Median z-score and combined p-values (using Fisher’s method) were then computed across different runs (at least two per sample). In order to be used in the logic formalisms, data were scaled between 0 (untreated control) and 1 (maximum activation). Conditions which were defined as not significantly different with respect to the untreated control (combined p-value < 0.05) were also set to 0.

For implementing and optimising the mathematical models, we used the CellNOptR tool (Terfve *et al*, 2012) and a modified version of the CNORode add-on to model logic-based ordinary differential equations (ODEs), as presented in (Eduati *et al*, 2017) and available at https://github.com/saezlab/CNORode2017. Using the CellNoptR package, the logic model was compressed as described in (Saez-Rodriguez *et al*, 2009) to reduce model complexity. In the logic ODE formalism (Wittmann *et al*, 2009), each node (i.e. species *x*_*i*_) is modeled by an ODE with a continuous update function *B*_*i*_ representing the regulation by the *N*_*i*_ upstream nodes.

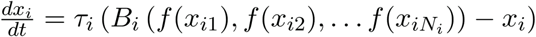

The tunable parameter *τ*_*i*_ represent the life-time of species *i*. We define each regulation using a sigmoidal transfer function:

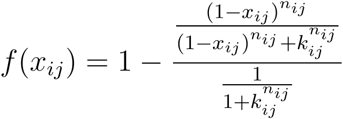

Where parameters *n* and *k* are fixed to 3 and 0.5 respectively, and the tunable parameter *k*_*ij*_ represent the strength of the regulation of species *j* on species *i* (edge *j*→*i*).

Parameters were estimated by fitting the model simulation to the experimental data using the optimization toolbox MEIGO (Egea *et al*, 2014). Bootstrapped distributions for all parameters were obtained by repeating the optimization resampling data with replacement.

### Statistical tests

Non-parametric tests were used because they are highly robust against non-normality. Pairwise comparisons (both on parameters and on predictions) for cell lines were performed using Wilcoxon rank sum test. Kruskal–Wallis rank sum test (one-way ANOVA on ranks) was used when comparing multiple groups (i.e. for patients, both on parameters and on predictions) and followed by post-hoc pairwise comparison with Wilcoxon sum rank test on the parameters which are not equally distributed among all groups. Effect size w was computed for Wilcoxon rank sum test as 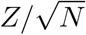, where *Z* is the statistics from the test and *√χ*^*2*^*/N* is the number of observations, and for the Kruskal–Wallis rank sum test it was computed as 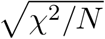 where *χ*^*2*^ is the statistics from the test and *N* is the number of observations. P-values were always Bonferroni adjusted to correct for multiple hypothesis testing. Significance thresholds (reported also in the main text) were set to to 0.01 for all adjusted p-values. For the effect size, the threshold was set to 0.2 when comparing model parameters and 0.3 when comparing predictions (to further limit the number of significant testable predictions).

### Validation experiments

AsPC1 and BxPC3 cells were used at passage 2-4 after thawing and seeded at 12,800 or 8,000 cells per well, respectively, in a 96-well plate with RPMI media (supplemented with 10% FBS, 1% penicillin/streptomycin, 4.5 mg/mL glucose and 1 mM sodium pyruvate) on day 0. Drugs were added on day 1 and viability was measured after 72 hours using CellTiter-Glo^®^ (Promega). Navitoclax has been used at fixed concentrations of 2.5 uM and 10 uM. Taselisib was used up to a concentration of 10 uM in an 8-points 1:3 dilution series, spanning a 2000-fold range. We performed three biological replicates. For each biological replicate each data point was measured in at least three wells per plate (i.e. technical replicates). Raw data were preprocessed by subtracting the average background (blank) and removing outliers only if one out of three technical replicate was off by >30% compared to the other two. This resulted in a maximum removal of two data points per plate. Viability data were normalised to the negative control condition (i.e. DMSO treated cells). We fitted a four-parameter log-logistic model using the ‘drm’ R package (Ritz *et al*, 2015) and computed the synergy score with Bliss independence model as implemented in the ‘synergyfinder’ R package (He *et al*, 2018).

## Acknowledgements

We thank Hayley Donnella, Jessica Wappler and Thorsten Cramer for useful discussion and feedback on the manuscript. We thank Diana Panayotova Dimitrova and Maria Feoktistova for constructive discussion about apoptosis pathway.

## Authors contributions

JSR and CM conceived the project. JSR and FE designed the project. FE performed the analysis under the supervision of JSR. PJ and MG performed experimental validation. FE and JSR wrote the manuscript. PJ contributed to manuscript finalization. All authors approved the final manuscript.

## References

Aksamitiene E, Kiyatkin A & Kholodenko BN (2012) Cross-talk between mitogenic Ras/MAPK and survival PI3K/Akt pathways: a fine balance. Biochem. Soc. Trans. 40: 139–146

Balmanno K & Cook SJ (2009) Tumour cell survival signalling by the ERK1/2 pathway. Cell Death Differ. 16: 368–377

Borisov N, Aksamitiene E, Kiyatkin A, Legewie S, Berkhout J, Maiwald T, Kaimachnikov NP, Timmer J, Hoek JB & Kholodenko BN (2009) Systems-level interactions between insulin–EGF networks amplify mitogenic signaling. Mol. Syst. Biol. 5: 256

Bykov VJN, Eriksson SE, Bianchi J & Wiman KG (2018) Targeting mutant p53 for efficient cancer therapy. Nat. Rev. Cancer 18: 89–102

Cagnol S & Chambard JC (2010) ERK and cell death: Mechanisms of ERK-induced cell death–apoptosis, autophagy and senescence. FEBS J. 277: 2–21

Eduati F, Doldàn-Martelli V, Klinger B & Cokelaer T (2017) Drug resistance mechanisms in colorectal cancer dissected with cell type–specific dynamic logic models. Cancer Res. 77: 3364–337

Eduati F, Utharala R, Madhavan D, Neumann UP, Longerich T, Cramer T, Saez-Rodriguez J & Merten CA (2018) A microfluidics platform for combinatorial drug screening on cancer biopsies. Nat. Commun. 9: 2434

Egea JA, Henriques D, Cokelaer T, Villaverde AF, MacNamara A, Danciu D-P, Banga JR & Saez-Rodriguez J (2014) MEIGO: an open-source software suite based on metaheuristics for global optimization in systems biology and bioinformatics. BMC Bioinformatics 15: 136

Fey D, Halasz M, Dreidax D, Kennedy SP, Hastings JF, Rauch N, Munoz AG, Pilkington R, Fischer M, Westermann F, Kolch W, Kholodenko BN & Croucher DR (2015) Signaling pathway models as biomarkers: Patient-specific simulations of JNK activity predict the survival of neuroblastoma patients. Sci. Signal. 8: ra130

Garcia-Alonso L, Iorio F, Matchan A, Fonseca N, Jaaks P, Peat G, Pignatelli M, Falcone F, Benes CH, Dunham I, Bignell G, McDade SS, Garnett MJ & Saez-Rodriguez J (2018) Transcription Factor Activities Enhance Markers of Drug Sensitivity in Cancer. Cancer Res. 78: 769–780

Grieco L, Calzone L, Bernard-Pierrot I, Radvanyi F, Kahn-Perlès B & Thieffry D (2013) Integrative modelling of the influence of MAPK network on cancer cell fate decision. PLoS Comput. Biol. 9: e1003286

He L, Kulesskiy E, Saarela J, Turunen L, Wennerberg K, Aittokallio T & Tang J (2018) Methods for High-throughput Drug Combination Screening and Synergy Scoring. Methods Mol. Biol. 1711: 351–398

Hill SM, Nesser NK, Johnson-Camacho K, Jeffress M, Johnson A, Boniface C, Spencer SEF, Lu Y, Heiser LM, Lawrence Y, Pande NT, Korkola JE, Gray JW, Mills GB, Mukherjee S & Spellman PT (2017) Context Specificity in Causal Signaling Networks Revealed by Phosphoprotein Profiling. Cell Syst 4: 73–83.e10

Iorio F, Knijnenburg TA, Vis DJ, Bignell GR, Menden MP, Schubert M, Aben N, Gonçalves E, Barthorpe S, Lightfoot H, Cokelaer T, Greninger P, van Dyk E, Chang H, de Silva H, Heyn H, Deng X, Egan RK, Liu Q, Mironenko T, et al (2016) A Landscape of Pharmacogenomic Interactions in Cancer. Cell 166: 740–754

Izetti P, Hautefeuille A, Abujamra AL, de Farias CB, Giacomazzi J, Alemar B, Lenz G, Roesler R, Schwartsmann G, Osvaldt AB, Hainaut P & Ashton-Prolla P (2014) PRIMA- 1, a mutant p53 reactivator, induces apoptosis and enhances chemotherapeutic cytotoxicity in pancreatic cancer cell lines. Invest. New Drugs 32: 783–794

Khoo KH, Hoe KK, Verma CS & Lane DP (2014) Drugging the p53 pathway: understanding the route to clinical efficacy. Nat. Rev. Drug Discov. 13: 217–236

Kim E, Kim J-Y, Smith MA, Haura EB & Anderson ARA (2018) Cell signaling heterogeneity is modulated by both cell-intrinsic and -extrinsic mechanisms: An integrated approach to understanding targeted therapy. PLoS Biol. 16: e2002930

King CC, Gardiner EM, Zenke FT, Bohl BP, Newton AC, Hemmings BA & Bokoch GM (2000) p21-activated kinase (PAK1) is phosphorylated and activated by 3-phosphoinositide-dependent kinase-1 (PDK1). J. Biol. Chem. 275: 41201–41209

Klinger B, Sieber A, Fritsche-Guenther R, Witzel F, Berry L, Schumacher D, Yan Y, Durek P, Merchant M, Schäfer R, Sers C & Blüthgen N (2013) Network quantification of EGFR signaling unveils potential for targeted combination therapy. Mol. Syst. Biol. 9: 673

Letai A (2017) Functional precision cancer medicine-moving beyond pure genomics. Nat. Med. 23: 1028–1035

Mai Z & Liu H (2009) Boolean network-based analysis of the apoptosis network: irreversible apoptosis and stable surviving. J. Theor. Biol. 259: 760–769

Matsuoka T & Yashiro M (2016) Molecular targets for the treatment of pancreatic cancer: Clinical and experimental studies. World J. Gastroenterol. 22: 776–789

Merkle R, Steiert B, Salopiata F, Depner S, Raue A, Iwamoto N, Schelker M, Hass H, Wäsch M, Böhm ME, Mücke O, Lipka DB, Plass C, Lehmann WD, Kreutz C, Timmer J, Schilling M & Klingmüller U (2016) Identification of Cell Type-Specific Differences in Erythropoietin Receptor Signaling in Primary Erythroid and Lung Cancer Cells. PLoS Comput. Biol. 12: e1005049

Morton JP, Timpson P, Karim SA, Ridgway RA, Athineos D, Doyle B, Jamieson NB, Oien KA, Lowy AM, Brunton VG, Frame MC, Evans TRJ & Sansom OJ (2010) Mutant p53 drives metastasis and overcomes growth arrest/senescence in pancreatic cancer. Proc. Natl. Acad. Sci. U. S. A. 107: 246–251

Ritz C, Baty F, Streibig JC & Gerhard D (2015) Dose-Response Analysis Using R. PLoS One 10: e0146021

Saez-Rodriguez J, Alexopoulos LG, Epperlein J, Samaga R, Lauffenburger DA, Klamt S & Sorger PK (2009) Discrete logic modelling as a means to link protein signalling networks with functional analysis of mammalian signal transduction. Mol. Syst. Biol. 5: 331

Saez-Rodriguez J, Alexopoulos LG, Zhang M, Morris MK, Lauffenburger DA & Sorger PK (2011) Comparing signaling networks between normal and transformed hepatocytes using discrete logical models. Cancer Res. 71: 5400–5411

Saez-Rodriguez J, MacNamara A & Cook S (2015) Modeling Signaling Networks to Advance New Cancer Therapies. Annu. Rev. Biomed. Eng. 17: 150814155959008

Schlatter R, Schmich K, Vizcarra IA, Scheurich P, Sauter T, Borner C, Ederer M, Merfort I & Sawodny O (2009) ON/OFF and Beyond - A Boolean Model of Apoptosis. PLoS Comput. Biol. 5: e1000595

She Q-B, Solit DB, Ye Q, O’Reilly KE, Lobo J & Rosen N (2005) The BAD protein integrates survival signaling by EGFR/MAPK and PI3K/Akt kinase pathways in PTEN-deficient tumor cells. Cancer Cell 8: 287–297

Tanaka H, Fujita N & Tsuruo T (2005) 3-Phosphoinositide-dependent Protein Kinase-1-mediated IκB Kinase β (IKKB) Phosphorylation Activates NF-κB Signaling. J. Biol. Chem. 280: 40965–40973

Tan N, Wong M, Nannini MA, Hong R, Lee LB, Price S, Williams K, Savy PP, Yue P, Sampath D, Settleman J, Fairbrother WJ & Belmont LD (2013) Bcl-2/Bcl-xL inhibition increases the efficacy of MEK inhibition alone and in combination with PI3 kinase inhibition in lung and pancreatic tumor models. Mol. Cancer Ther. 12: 853–864

Terfve C, Cokelaer T, Henriques D, MacNamara A, Goncalves E, Morris MK, van Iersel M, Lauffenburger DA & Saez-Rodriguez J (2012) CellNOptR: a flexible toolkit to train protein signaling networks to data using multiple logic formalisms. BMC Syst. Biol. 6: 133

Türei D, Korcsmáros T & Saez-Rodriguez J (2016) OmniPath: guidelines and gateway for literature-curated signaling pathway resources. Nat. Methods 13: 966–967

Wang L, Du F & Wang X (2008) TNF-alpha induces two distinct caspase-8 activation pathways. Cell 133: 693–703

Werner HMJ, Mills GB & Ram PT (2014) Cancer Systems Biology: a peek into the future of patient care? Nat. Rev. Clin. Oncol. 11: 167–176

Wittmann DM, Krumsiek J, Saez-Rodriguez J, Lauffenburger DA, Klamt S & Theis FJ (2009) Transforming Boolean models to continuous models: methodology and application to T-cell receptor signaling. BMC Syst. Biol. 3: 98

Wörmann SM, Song L, Ai J, Diakopoulos KN, Kurkowski MU, Görgülü K, Ruess D, Campbell A, Doglioni C, Jodrell D, Neesse A, Demir IE, Karpathaki A-P, Barenboim M, Hagemann T, Rose-John S, Sansom O, Schmid RM, Protti MP, Lesina M, et al (2016) Loss of P53 Function Activates JAK2-STAT3 Signaling to Promote Pancreatic Tumor Growth, Stroma Modification, and Gemcitabine Resistance in Mice and Is Associated With Patient Survival. Gastroenterology 151: 180–193.e12

Yaffe MB (2013) The Scientific Drunk and the Lamppost: Massive Sequencing Efforts in Cancer Discovery and Treatment. Sci. Signal. 6: scisignal.2003684v1

Zañudo JGT, Steinway SN & Albert R (2018) Discrete dynamic network modeling of oncogenic signaling: Mechanistic insights for personalized treatment of cancer. Current Opinion in Systems Biology 9: 2452–3100

